# The conservation of a core virome in *Aedes* mosquitoes across different developmental stages and continents

**DOI:** 10.1101/2020.04.23.058701

**Authors:** Chenyan Shi, Lu Zhao, Evans Atoni, Weifeng Zeng, Xiaomin Hu, Jelle Matthijnssens, Zhiming Yuan, Han Xia

## Abstract

Mosquitoes belonging to the genus *Aedes* can efficiently transmit many pathogenic arboviruses, placing a great burden on public health worldwide. In addition, they also carry a number of insect specific viruses (ISVs), and it was recently suggested that some of these ISVs might form a stable species-specific “core virome” in mosquito populations. However, little is known about such a core virome in laboratory colonies and if it is present across different developmental stages. In this study, we compared the viromes in eggs, larvae, pupae and adults of *Aedes albopictus* mosquitoes collected from the field as well as from a lab colony. The virome in lab-derived *Ae. albopictus* is very stable across all stages, consistent with a vertical transmission route of these viruses, forming a “vertically transmitted core virome”. The different stages of field collected *Ae. albopictus* mosquitoes also contains this stable vertically transmitted core virome as well as another set of viruses shared by mosquitoes across different stages, which might be an “environment derived core virome”. Both these vertically and environmentally transmitted core viromes in *Ae. albopictus* deserve more attention with respect to their effects on vector competence for important medically relevant arboviruses. To further study this core set of ISVs, we screened 46 publically available SRA viral metagenomic dataset of mosquitoes belonging to the genus *Aedes*. Some of the identified core ISVs are identified in the majority of SRAs. In addition, a novel virus, Aedes phasmavirus, is found to be distantly related to Yongsan bunyavirus 1, and the genomes of the core virus Phasi Charoen-like phasivirus is highly prevalent in the majority of the tested samples, with nucleotide identities ranging from 94% to 99%. Finally, Guadeloupe mosquito virus, and some related viruses formed three separated phylogenetic clades. How these core ISVs influence the biology of mosquito host, arboviruses infection and evolution deserve to be further explored.

## 1. Introduction

Mosquitoes are highly diverse and widely disseminated. They occupy numerous biotopes and are potential vectors for several pathogenic arboviruses. Specifically*, Aedes* sp. impose a great threat to global public health. This is due to their ecological plasticity traits that include egg diapause, versatility in using natural and/or urban breeding spots [1] and opportunistic feeding patterns [2, 3], which might have promoted their dispersion and adaptation to new uncolonized territories that range from tropical to temperate regions [4].

Moreover, a large number of pathogenic arboviruses, such as Dengue virus (DENV), Yellow fever virus (YFV), Zika virus (ZIKV) and Chikungunya virus (CHIKV), are efficiently transmitted by *Aedes* (*Ae.*) mosquitoes, especially *Ae. aegypti* and *Ae. Albopictus* [5]. DENV and CHIKV are the most common arboviral diseases, with more than 40% of the world’s population living in areas with transmission of either of these viruses [6, 7]. Particularly, DENV epidemics are becoming a major public health concern in Guangdong Province of China. It experienced an unexpectedly explosive outbreak in 2014 with 45,224 reported dengue fever cases [8], which is more than the total number of cases in the past 30 years in this region [9]. YFV remains endemic in tropical regions in Africa as well as South and Central America, and despite the presence of a good vaccine and large vaccination campaigns in the past, Angola and neighbouring Democratic Republic of the Congo experienced a large outbreak in recent years [10]. ZIKV is the most recent emergent arbovirus, receiving attention worldwide due to its explosive spread in South America, and its association with foetal microcephaly and Guillain-Barré syndrome [11, 12].

Due to the lack of therapeutic treatments and widely available vaccines, control of mosquito population is the most efficient way to prevent arboviral diseases [13]. Mosquitoes are holometabolous insects, whose life cycle goes through four separate stages, including eggs, larva, and pupa in an aquatic habitat and a subaerial adult stage. Although only the females with anthropophilic blood feeding habit can transmit arboviruses, reducing the population of mosquito on the juvenile stages will decrease the number of adults and subsequently lower the transmission chances. The conventional control strategies, including the use of insecticide and environmental sanitation, are facing challenges due to their sustainability and organizational complexity [13, 14]. In light of the holobiont concept, mosquito-associated microbial communities play an important role in host biology, which may provide new strategies for mosquito control [15].

To date, most of the conducted mosquito microbiota studies have solely focused on the bacterial component in mosquitoes and their possible intrinsic roles in mosquito biology [16]. For example, the nutrient produced by symbiotic bacteria is very important for larvae development [17], and the endosymbiotic alpha-proteobacteria *Wolbachia* is a natural endosymbiont in several mosquito species and can induce cytoplasmic incompatibility [18]. When this bacterium is introduced into its not-natural host *Ae. aegypti*, it has strong suppressive effect on DENV infection [19, 20]. More interestingly, some newly discovered mosquito specific viruses have been proposed as future biological control agents against arboviruses [21–23], as novel vaccine platforms [23] or used in diagnostic assays in chimeric virus formation with structural protein genes from arboviruses [23], However, little is known about the virome community dynamics in mosquitoes during its holometabolous development. Some studies have performed viral metagenomics on field mosquitoes collected from different countries (including the United States, Puerto Rico, China, Kenya, Australia, Sweden, etc.), mainly focusing on novel virus discovery [24–32]. In a recent study we showed the presence of a relatively stable “core eukaryotic virome” in field *Ae. aegypti* and *Culex quinquefasciatus* mosquitoes from Guadeloupe [33]. However, the presence of insect specific viruses (ISVs) in mosquito lab colonies is barely known, which limit our understanding in their role in the development and physiology of mosquitoes. On the other hand, knowing the normal healthy background virome as a reference, will allow us to distinguish between inherent vertically transmitted components and components acquired from the environment of the field mosquitoes, which will benefit the identification of potential viral pathogens and improve the reliability of risk assessment using lab mosquitoes.

The aim of this study was to analyze the core virome structure of *Ae. albopictus* mosquitoes during distinct developmental stages, in a laboratory-derived colony originally obtained from Jiangsu Province, China, and to compare these findings to the virome of field *Ae. albopictus* mosquitoes collected in Guangzhou (Guangdong Province, China), around 1300 kilometer apart. Furthermore, we conducted a comparative meta-analysis between the viruses identified in this study and 46 publicly available virome SRA datasets of *Aedes* mosquitoes. Finally, phylogenetic analyses were performed on three selected viruses: Aedes phasmavirus which is highly present in both field and lab *Ae. albopictus* from China; Guadeloupe mosquito related virus and Phasi Charoen-like phasivirus, which were found in the majority of investigated virome data sets.

## 2. Materials and methods

### 2.1 Sample collection

Field larva, pupa and adult samples of *Ae. albopictus* were trapped in Guangzhou (Liwan District), Guangdong Province of China in July-November of 2017 and 2018. In Kenya, adult mosquitoes of *Ae. aegypti* were trapped during July and August of the year 2018 in Ukunda and Kisumu. All samples were collected from residential quarter in urban area, transported to the laboratory using an appropriate cold chain and stored at −80 °C. Mosquito species were determined by morphological identification and samples were assigned into pools according to date of collection (Table 1). An *Ae. albopictus* colony was established in our laboratory in 2017 (Wuhan, China), which originated from the stable colony from the National Institute for Communicable Disease Control and Prevention, China CDC (Beijing, China). The adult mosquitoes were held at (27–30 °C), with 60–85% relative humidity, and a photoperiod of 12:12 h light-dark cycle. The larvae were fed with a mixture of yeast powder and wheat mill. Adult mosquitoes were placed into cages (30 cm × 30 cm×30 cm) and the males were fed with 10% sucrose solution, while the females were fed with blood from mice. The egg, larva, pupa, male and female adult samples of lab colonies mosquitoes were collected respectively in August of 2017 (Table 1).

**Table 1.**
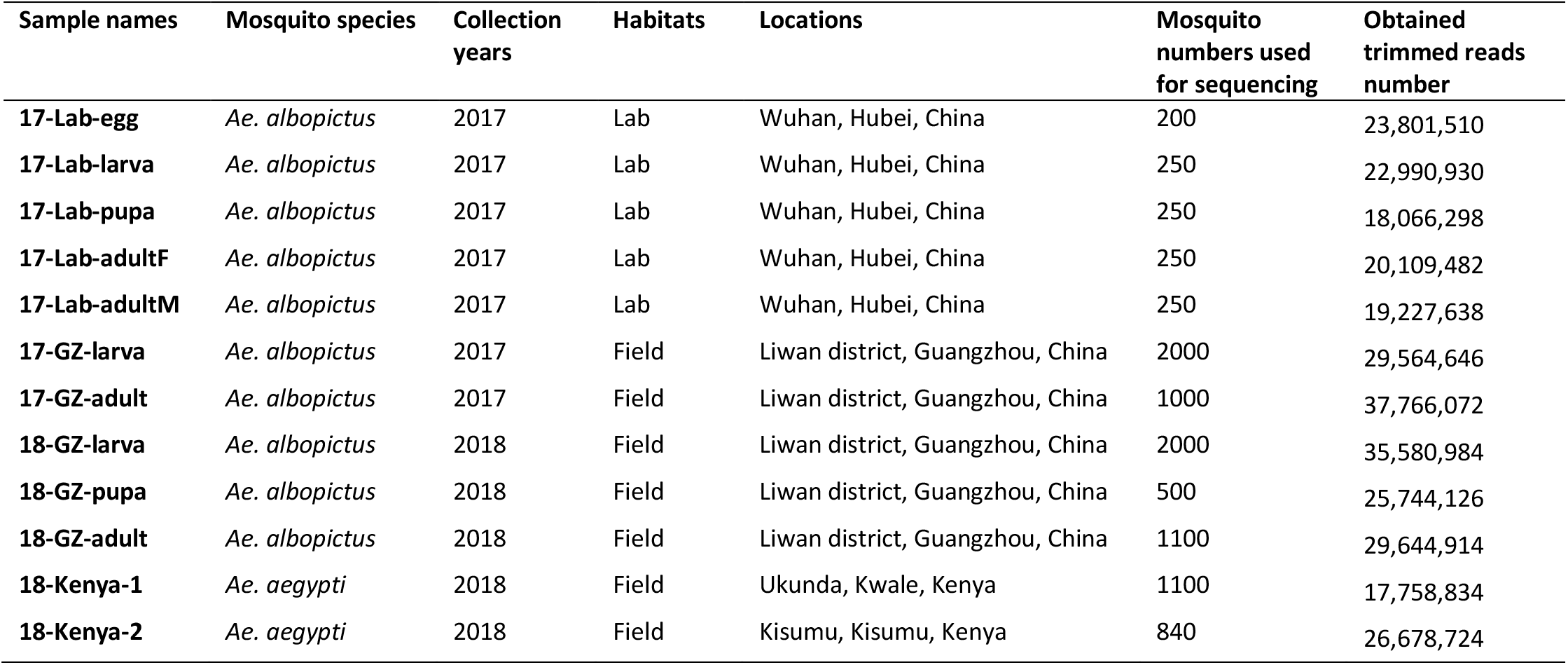
*Aedes* spp. from China and Kenya used for viral metagenomic sequencing.

| Sample names | Mosquito species | Collection years | Habitats | Locations | Mosquito numbers used for sequencing | Obtained trimmed reads number |
| --- | --- | --- | --- | --- | --- | --- |
| 17-Lab-egg | *Ae. albopictus* | 2017 | Lab | Wuhan, Hubei, China | 200 | 23,801,510 |
| 17-Lab-larva | *Ae. albopictus* | 2017 | Lab | Wuhan, Hubei, China | 250 | 22,990,930 |
| 17-Lab-pupa | *Ae. albopictus* | 2017 | Lab | Wuhan, Hubei, China | 250 | 18,066,298 |
| 17-Lab-adultF | *Ae. albopictus* | 2017 | Lab | Wuhan, Hubei, China | 250 | 20,109,482 |
| 17-Lab-adultM | *Ae. albopictus* | 2017 | Lab | Wuhan, Hubei, China | 250 | 19,227,638 |
| 17-GZ-larva | *Ae. albopictus* | 2017 | Field | Liwan district, Guangzhou, China | 2000 | 29,564,646 |
| 17-GZ-adult | *Ae. albopictus* | 2017 | Field | Liwan district, Guangzhou, China | 1000 | 37,766,072 |
| 18-GZ-larva | *Ae. albopictus* | 2018 | Field | Liwan district, Guangzhou, China | 2000 | 35,580,984 |
| 18-GZ-pupa | *Ae. albopictus* | 2018 | Field | Liwan district, Guangzhou, China | 500 | 25,744,126 |
| 18-GZ-adult | *Ae. albopictus* | 2018 | Field | Liwan district, Guangzhou, China | 1100 | 29,644,914 |
| 18-Kenya-1 | *Ae. aegypti* | 2018 | Field | Ukunda, Kwale, Kenya | 1100 | 17,758,834 |
| 18-Kenya-2 | *Ae. aegypti* | 2018 | Field | Kisumu, Kisumu, Kenya | 840 | 26,678,724 |

### 2.2 Sample preparation for NGS

Pooled mosquito samples were triturated by a Tgrinder electric tissue grinder OSE-Y30 (TIANGEN,China) on ice using sterile pestles with 200 μl of Roswell Park Memorial Institute (RPMI) medium. Mosquito homogenates were clarified by centrifugation at 20,000×g (4 °C for 30 min) and filtered through a 0.45 μm membrane filter (Millipore, Billerica, USA). Supernatants were collected and stored at −80 °C. RNA was extracted from the supernatants per the manufacturer’s instructions by using RNeasy mini kit (Qiagen, Germany). Then strand-specific libraries were prepared using the TruSeq® Stranded Total RNA Sample Preparation kit (Illumina, USA). TruSeq PE (paired-end) Cluster Kit v3 (Illumina, USA) and TruSeq SBS Kit v3 - HS (300 Cycles) (Illumina, USA) were used for sequencing with 150 bp per read (PE 2×150 bp), which was performed at Shanghai Biotechnology Corporation with the Illumina HiSeq 2500 platform (Illumina, USA).

### 2.3 Bioinformatic analysis

The obtained raw paired-end reads were trimmed for quality and adapters using Trimmomatic [34] and the remaining reads were *de novo* assembled into contigs using metaSPAdes [35]. Contigs from all pools longer than 500bp were filtered for redundancy at 95% nucleotide identity over 80% of the length using ClusterGenomes [36]. The representative contigs collection was classified using DIAMOND [37] against the nr database (updated on 29th Sep 2019) on sensitive mode for taxonomic annotation. KronaTools [38] were used to parse the output file of DIAMOND, which found the least common ancestor of the best 25 DIAMOND hits (based on BLASTx score) for each contig. All contigs annotated as eukaryotic virus were extracted using an in-house python script. The abundance and length coverage of eukaryotic virus contigs in individual pool were obtained by aligning trimmed and decontaminated reads of each samples to the representative contigs collection using BBMap [39]. Abundance tables from eukaryotic virus were extracted and further used for making heatmaps in R with the ComplexHeatmap [40], ggplot2 [41], phyloseq [42] packages. The length coverage of each viral species per sample was calculated by dividing the contigs length covered (by at least one read) by the total length of the contigs.

### 2.4 Eukaryotic virus screening of *Aedes* mosquito virome data in SRA

To investigate the level of conservation of the eukaryotic core viruses identified in Chinese samples from this study with those of other *Aedes* mosquitoes worldwide, we retrieved 46 published SRA datasets (including 25 datasets of *Aedes* sp. from Guadeloupe [33], eight from Puerto Rico [24], four from USA [24], seven from Australia [43, 44], one from Thailand [44] and one from China [45]) (Additional file 1 and Fig. 1). Two more viral metagenomic data of *Ae. aegypti* collected in Kenya were also used (Table 1 and Fig. 1). The world map displaying the geographic origin of all *Aedes* virome datasets used in this study were drawn with ArcGIS (ArcMap 10.5). The trimmed reads of the two datasets of *Ae. aegypti* from Kenya were *de novo* assembled using metaSPAdes [35]. The raw reads of SRA data sets were *de novo* assembled by SKESA [46] with default settings. These obtained contigs were taxonomically annotated using DIAMOND against the nr database (updated on 29th Sep 2019) on sensitive mode. KronaTools [38] and an in-house python script were used to parse the output file of DIAMOND and extracted eukaryotic virus contigs.

**Fig 1.**
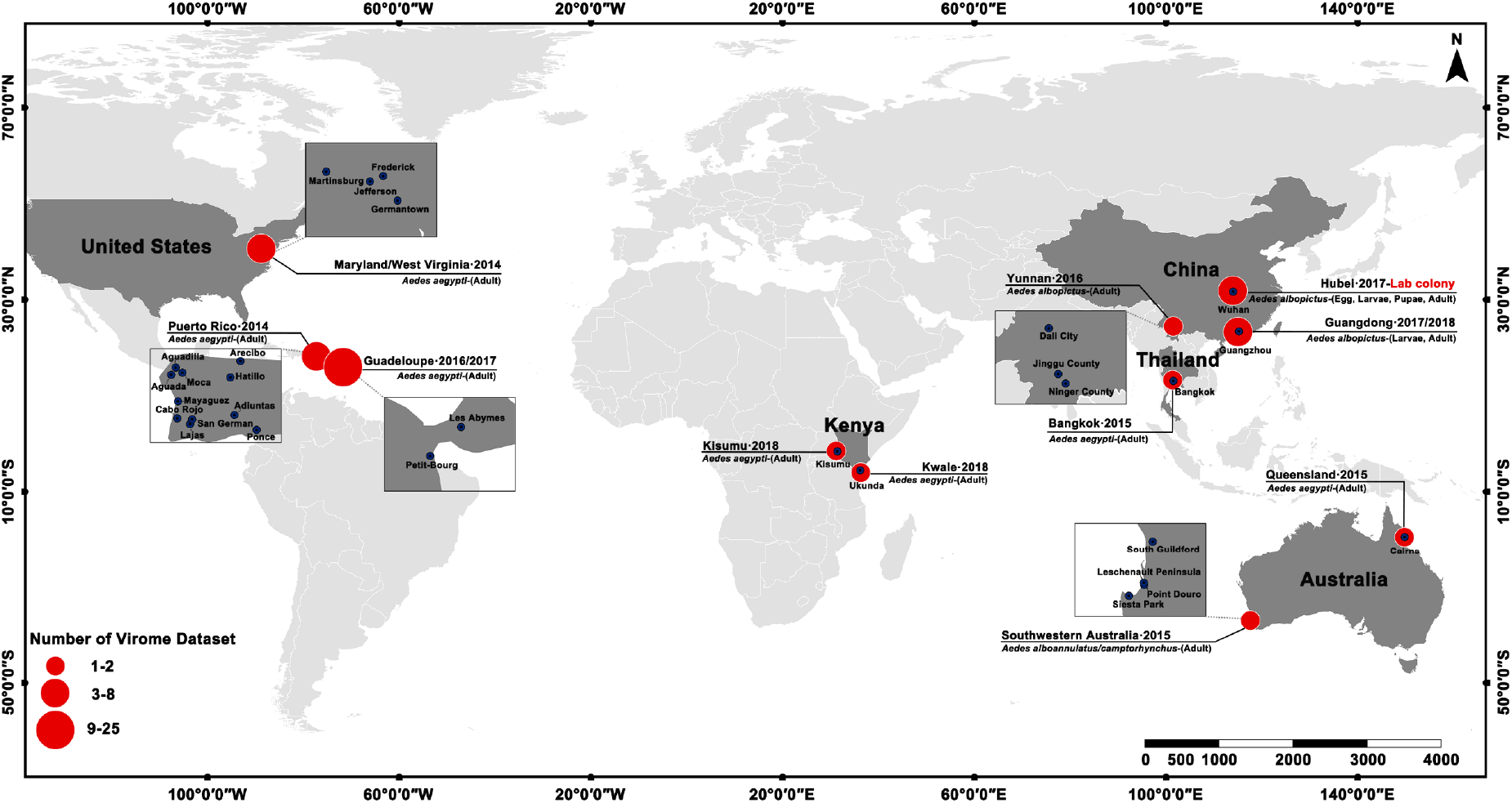
Global distribution of *Aedes* mosquitoes virome datasets used in this study.

### 2.5 The presence of eukaryotic virus in all *Aedes* virome datasets

Eukaryotic virus contigs longer than 500 bp from all 58 Aedes virome datasets were merged according to their taxonomical annotation. A viral species was considered as presence in the sample as long as the sample had one contig (>500 bp) assigned to the species. The viral species are shown in the heatmap (Figure 3) only if they were present in samples from at least three countries and contain at least one contig that was longer than 1500bp.

### 2.6 Phylogenetic analysis

To obtain the complete genomes related to Yongsan bunyavirus 1 (YBV1), Phasi Charoen-like phasivirus (PCLPV) and Guadeloupe mosquito virus (GMV) from all *Aedes* virome datasets, the trimmed reads of each sample were individually mapped (with BWA [47]) against the selected reference genomes: 17-Lab-egg-L (MT361040), 17-Lab-egg-M (MT361040) and 17-Lab-pupa-S (MT361044) as the reference for YBV1, PCLPV isolate Rio (KR003786.1, KR003784.1 and KR003785.1) for PCLPV and 18-GZ-larva-seg1 (MT361057) and 18-GZ-pupa-seg2 (MT361060) for GMV. The consensus sequences of these viruses were generated from the bam files using samtools and bcftools [48]. Accession numbers of these viruses obtained from our dataset are in Additional file 2 and viral genome sequences recovered from SRA datasets can be found in https://github.com/Matthijnssenslab/AedesVirome.

The nucleotide sequence of the complete genome or complete coding region of each genome (segment) of these viruses were used to determine the evolutionary history of the discovered viruses together with appropriate reference strains from GenBank. Alignments of sequences were performed with MAFFT v7.222 [49] using the most accurate algorithm L-INS-I with 1000 cycles of iterative refinement. Ambiguously aligned regions were removed by trimAl v1.2 [50] using automated trimming heuristic, which is optimized for Maximum-Likelihood (ML) phylogenetic tree reconstruction. Phylogenetic trees for each segment were reconstructed from 1,000 ultrafast bootstrap maximum likelihood (ML) tree replicates using IQ-TREE v1.6.11 [51] with best-fit model selection by ModelFinder [52]. FigTree v1.4.3 [53] was used for phylogenetic trees visualization.

## 3 Results

### 3.1 Comparison of the eukaryotic virome in field and lab *Ae. albopictus*

The egg, larva, pupa and adult (male and female) pools from a lab colony in Wuhan, as well as the larva, pupa and adult pools from the field in Guangzhou underwent the metagenomic sequencing and an average of 27 million trimmed reads were obtained per pool (Table 1), which were *de novo* assembled into 1,657,229 contigs in total. The clustering of 71,303 contigs longer than 500 bp from all samples at 95% nucleotide identity over 80% of the length results in 56,419 representative contigs. According to BLASTx annotation results using Diamond, the majority of the representative contigs (93%) belonged to Eukaryota (mainly derived from the mosquito host genome) and 179 contigs were assigned as eukaryotic viruses. Eukaryotic viral reads in each pool occupied 0.2% to 1.8% of trimmed reads, except for pool 17-GZ-larva which contained 15.8% viral reads. Most of the eukaryotic viral contigs were annotated as belonging to 80 different viruses (including several viruses with segmented genomes), although some of these had very low similarities to known viruses in the database (Fig. 2). No known pathogenic arboviruses were identified, and the closest relatives of the identified viruses were generally found in insects or plant. Only 20 of the 80 viral species belonged to an established viral genus or family (e.g. *Flavivirus*, *Iflavirus*, *Phasivirus*, *Quaranjavirus*, *Rhabdoviridae*, *Virgaviridae*, *Totiviridae*, *Nodaviridae*), whereas the remaining ones are currently unclassified.

**Fig. 2.**
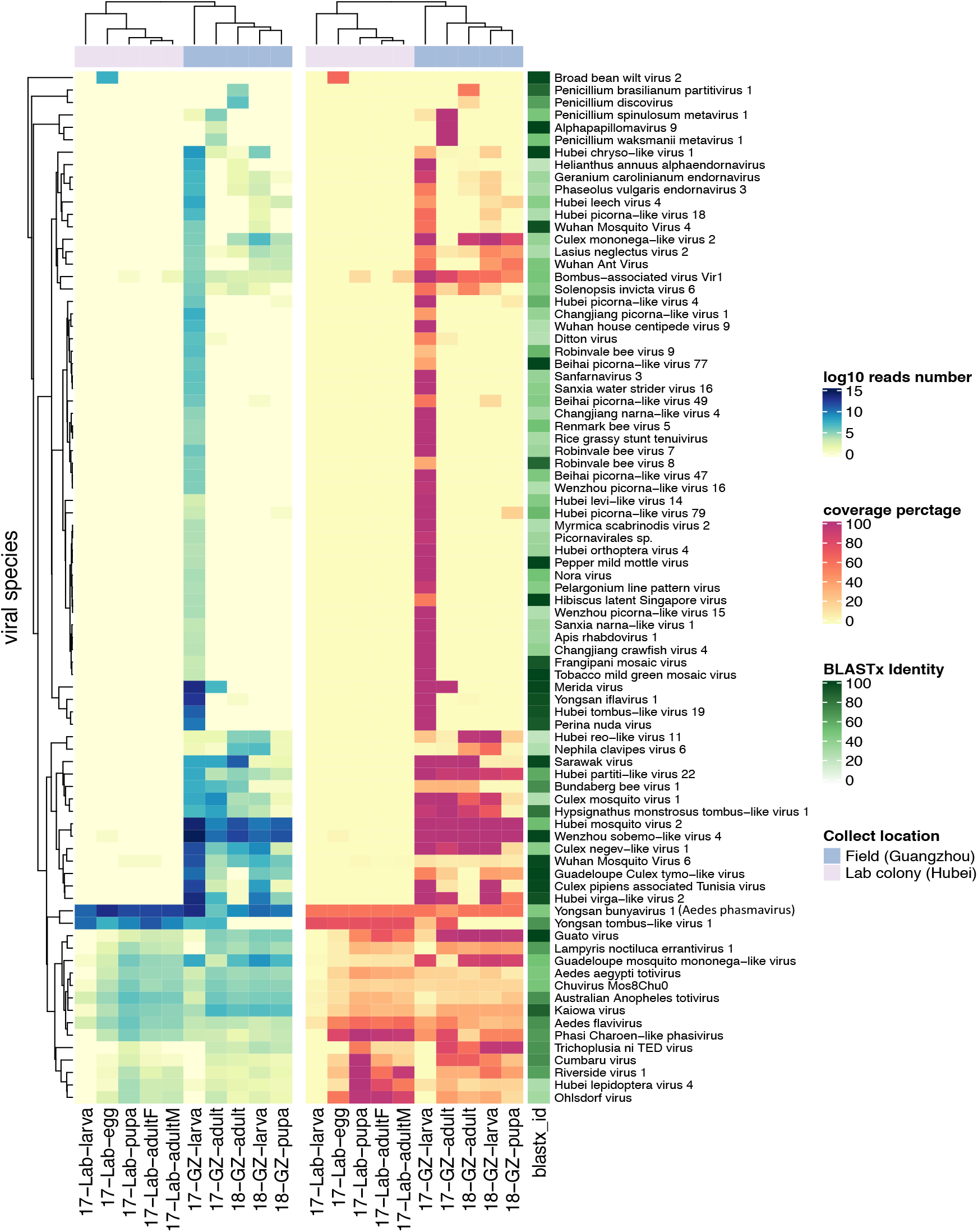
Abundance and length coverage of eukaryotic viral species in wild and lab-derived *Aedes albopictus* pools. The heatmaps show the total mapped reads number on a log10 scale (left) and length coverage (right) of assembled contigs of each virus. The hierarchical clustering is based on the Bray–Curtis distance matrix calculated from the log10 reads number. The viral species names shown in the heatmap are from the taxonomic annotation by DIAMOND and KronaTools. For each of the contigs assigned to a particular species, the ORF with the highest BLASTx identity to a reference sequence was taken, and the median identity of these different ORFs is shown in the shaded green bar.

The heatmap in Fig. 2 displays the mapped read numbers against each of the representative (partial) viral genome as well as the length coverage of assembled contigs of each virus (Additional File 3 and 4). The virome in field *Ae. albopictus* collected from both 2017 and 2018 showed significantly higher richness and diversity compared to that in lab-derived mosquito pools. The adjusted p-value of Wilcoxon test on Chao1 and Shannon index were 0.012 and 0.008, respectively (Additional File 5). The virome profile of the different pools of lab colony are relatively stable, except that several viral species are absent or low abundant in the larvae pool (Fig. 2). The wild larvae, pupa and adult pools collected in GZ in 2018 and the adult pool in 2017 also display a very similar viral community. However, the larvae pool collected in GZ 2017 contained more than 30 additional unique viral species with almost 100% length coverage, which are almost completely absent in the adults collected at the same time and in the same place. Furthermore, the lab and field *Ae. albopictus* have 20 viral species in common. Among them, some contigs only distantly related (47% BLASTx identity) to YBV1, are present in all lab and field derived mosquito pools with a high abundance and length coverage. This is also true for Phasi Charoen-like phasivirus, and contigs annotated as Yongsan tombus-like virus 1, except in field mosquitoes collected in 2018. Some viruses such as Phasi Charoen-like phasivirus appear to be present in in higher levels in the lab-derived *Ae. albopictus* samples. In contrast, reads of some viruses appear to be present in higher number in the field samples, as is the case for Guato virus and Guadeloupe mosquito mononega-like virus. Furthermore, some viruses are only present in field mosquitoes among all pools with high abundance, such as Hubei partiti—like virus 22, Hubei mosquito virus 2 and Wenzhou sobemo−like virus 4.

### 3.2 Prevalence of viruses in public SRA data sets derived from *Aedes* mosquitoes

All 46 available viral metagenomic data of *Aedes* sp. from public database (SRA) together with two samples from Kenya (Additional file 1) were further analyzed to determine the prevalence of conserved ISVs in various *Aedes* mosquitoes. These samples were collected from locations in different continents including the USA, Puerto Rico, Australia, Thailand, Guadeloupe, China and Kenya. The mosquito species in the majority of samples are *Ae. aegypti*, except the one from China (*Ae. Albopictus*) and five from Southwestern Australia (*Ae. alboannulatus* or *Ae. Camptorhynchus*). The heatmap in Fig. 3 shows the presence and absence of 42 viruses in the 58 analyzed virome data sets, grouped per collection locations and viral families. Only the viral species present in minimum three locations and containing at least one contig longer than 1.5 kb are displayed in the figure. Notably, the total number of viral species present in samples from the USA and Puerto Rico are much less than in other locations, which may due to the low sequencing depths, , or different viruses present in these samples.

**Fig. 3.**
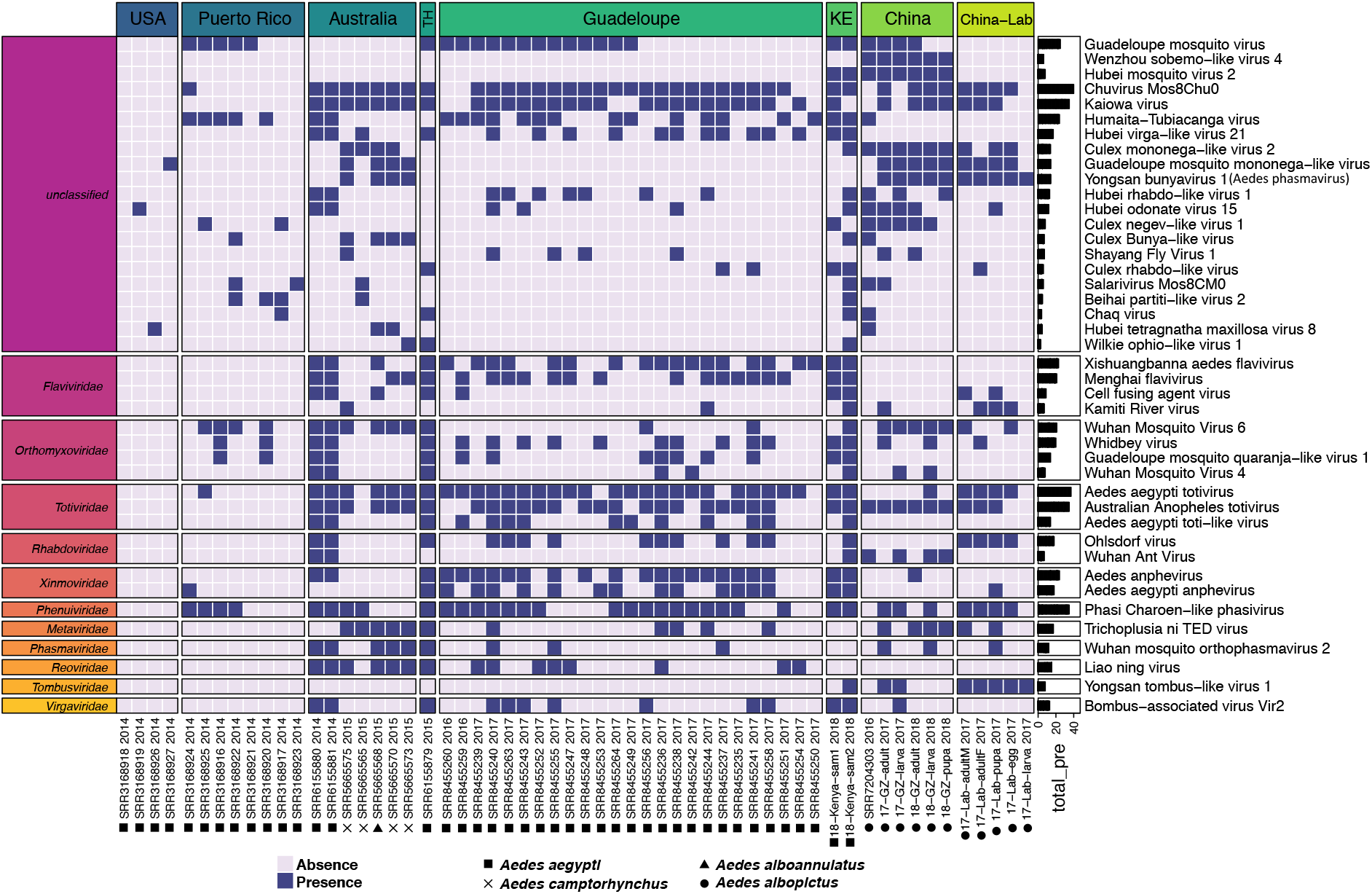
Presence and Absence of viral species in *Aedes* sp. virome datasets. The viral species shown in the heatmap present in samples from at least three locations and contain at least one contig that is longer than 1500 bp. The specific viral species was considered as presence in the sample, as long as the sample had one contigs (>500 bp) assigned to the species.

*Totiviridae* is the most prevalent viral family, containing two highly abundant species (Aedes aegypti totivirus and Australian Anopheles totivirus) with positivity rates of 63.8% and 60.3%, respectively. Phasi Charoen-like phasivirus in the *Phenuiviridae* is present in all four *Aedes* species and all locations (except USA, which only has three viruses presented). In addition, the families *Flaviviridae*, *Orthomyxoviridae*, *Rhabdoviridae* and *Xinmoviridae* also contain viral species present in 5 or more locations and at least *Ae. aegypti* and *Ae. albopictus* mosquitoes. For the unclassified viral species, Chuvirus Mos8Chu0 and Kaiowa virus are present in 40 and 35 out of 58 samples, respectively, and were found in all four *Aedes* species. Guadeloupe mosquito virus is detected in field *Ae. aegypti* and *Ae. albopictus mosquitoes* from Puerto Rico, Thailand, Guadeloupe, Kenya and China. Humaita-Tubiacanga virus (absent in lab-derived and wild *Ae. albopictus* from this study) are found in field *Ae. albopictus* from Yunnan (China) and field *Ae. aegypti* from 5 locations. Some of the viruses that are highly prevalent in both lab and field *Ae. albopictus* from China, including Culex mononega-like virus 2, Guadeloupe mosquito mononega-like virus and YBV1, are absent in most countries except in *Ae. camptorhynchus* from Australia.

### 3.3 Phylogeny of three selected viruses

Phylogenetic analysis was conducted on three selected viruses related to YBV1, which presented in both field and lab *Ae. albopictus* from China with high abundance, as well as the highly prevalent PCLPV and newly described GMV. Recovered coding complete viral genomes related to YBV1, PCLPV and GMV from analyzed viral metagenomic data of *Aedes* mosquitoes as well as corresponding reference genomes from GenBank were used for phylogeny. YBV1 is a newly identified virus reported in 2019, which has a high prevalence in *Ae. vexans nipponii* from Korea [54], and classified in the family *Phasmaviridae*. A novel virus of this family (tentatively named as Aedes phasmavirus in this study) distantly related to YBV1 (*vide infra*) was presented in lab *Ae. albopictus* from China and filed *Aedes* mosquitoes collected from three countries, including China, Korea, and Australia (Table 2). PCLPV, a widely distributed mosquito viruses, has been identified from lab colonies, mosquito cell lines and wild *Aedes* mosquitoes in eight counties or regions (China, Puerto Rico, Thailand, Kenya, USA, Australia, Guadeloupe, Brazil) (Table 2). It has been suggested that is maintained in nature through vertical transmission [55]. A recent study has reported that GMV is closely related to Wenzhou sobemo-like virus 4 and Hubei mosquito virus 2 [56]. GMV related virus has been detected only in mosquito sample collected from the field, indicating it is likely acquired horizontally from the environment.

**Table 2.** The distribution of three selected viruses.

|  | region | origin | Aedes<br>phasmavirus<br>(relate to YBV1) | PCLPV | GMV related virus |
| --- | --- | --- | --- | --- | --- |
| lab | China | 2017-Wuhan-lab | + | + |  |
|  | Belgium | 2018-Belgium-lab |  | + |  |
|  | UK | 2017-Bristol-lab |  | + |  |
| field | China | 2016-Zhejiang |  | + | + |
|  |  | 2017-Guangzhou | + | + | + |
|  |  | 2018-Guanghozu | + | + | + |
|  | Puerto Rico | 2014-Puerto Rico |  | + | + |
|  | Thailand | 2008-Thailand |  | + |  |
|  |  | 2015-Thailand |  | + | + |
|  | Kenya | 2018-Kenya1 |  | + | + |
|  |  | 2018-Kenya2 |  | + | + |
|  | USA | 2014-USA |  |  |  |
|  |  | 2015-USA |  | + |  |
|  | Australia | 2014-Australia |  | + |  |
|  |  | 2015-Australia | + | + |  |
|  |  | 2016-Australia |  | + |  |
|  | Korea | 2016-Korea | + |  |  |
|  | Guadeloupe | 2016-Guadeloupe |  | + | + |
|  |  | 2017-Guadeloupe |  | + | + |
|  | Brazil | 2012-Brazil |  | + |  |

#### 3.3.1 Phylogeny of Aedes phasmavirus

In both lab and field derived *Ae. albopictus* sequence data from China in 2017 and 2018, three contigs of Aedes phasmavirus (APV) in each sample were found to share highest amino acid identities of 50.8%, 48.1%, and 43.6% with S-, M- and L-segments of YBV1 (MH703047.1, MH703046.1, MH703045.1), respectively. In total, 10 viral genomes of APV with three segments could been recovered from all lab and field *Ae. albopictus* pools of this study in China (17-lab-egg, 17-lab-larva, 17-lab-pupa, 17-lab-adultM, 17-lab-adultF, 17-GZ-larva, 17-GZ-adult, 18-GZ-larva, 18-GZ-pupa, 18-GZ-adult) with the longest lengths of 1185, 2022, and 6468 nt corresponding for the complete cds (coding sequence) of the nucleocapsid, glycoprotein and RNA-dependent RNA polymerase (RdRp) genes, respectively. For all three segments, the nucleotide identity among the 10 assembled viral genomes ranged between 96% and 100%, which indicated they represent closely related variants. In the phylogenetic analysis of the three segments separately, the 10 viral genomes of APV clustered within the genus *Orthophasmavirus* in the family of *Phasmaviridae* (Fig. 4). They closest relative was YBV1 identified in *Ae. vexans* mosquitoes collected in South Korea in 2016, but with low nucleotide similarities (ranging from 52% to 57%) in the coding regions of the S-, M-, and L-segments. Thus, the APVs presented in the lab and field *Ae. albopictus in* China appears to represent a novel species in the genus *Orthophasmavirus*.

**Fig. 4.**
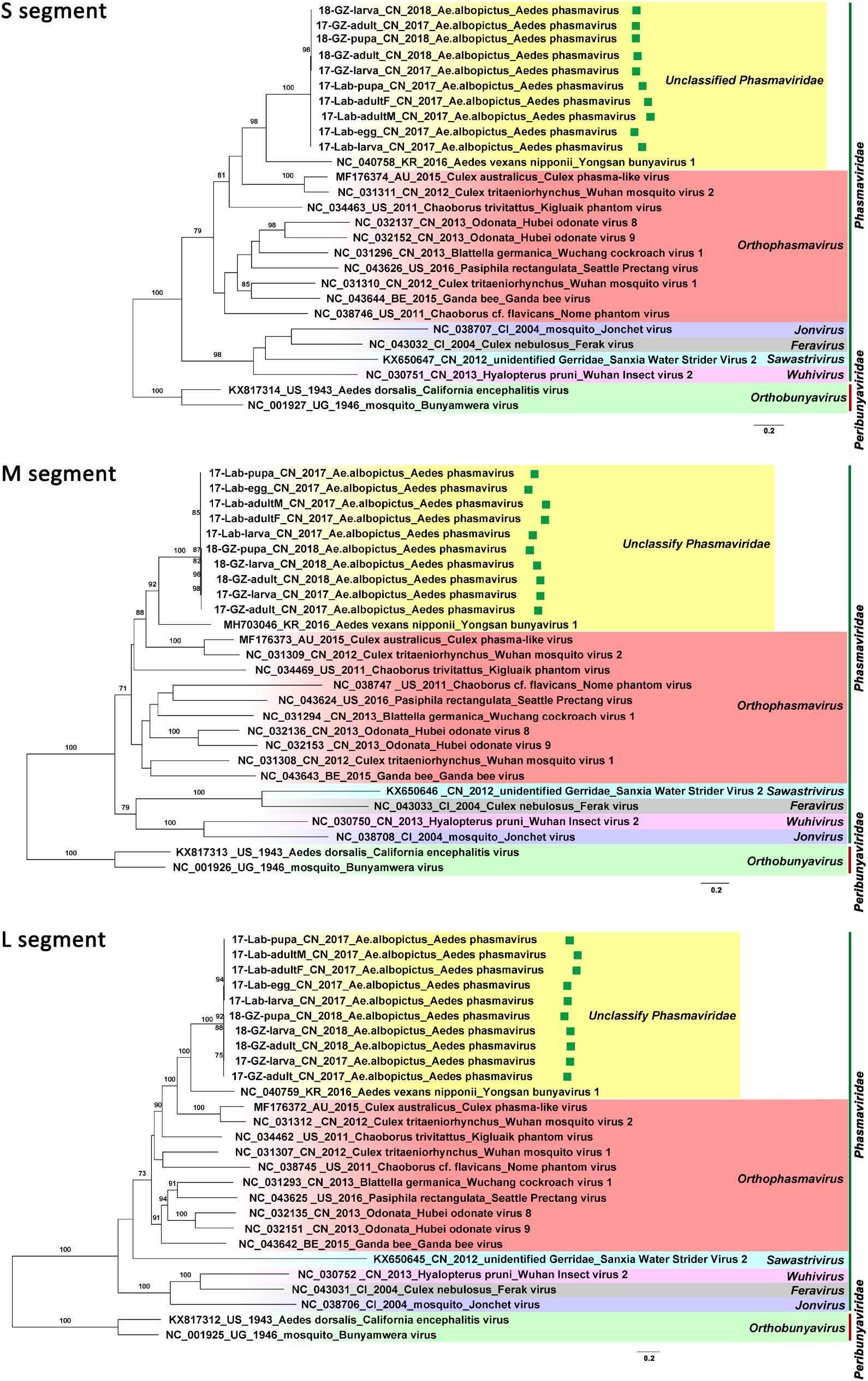
Maximum likelihood phylogeny on the nucleotide level of complete cds region in S, M, and L segment of the newly found *Orthophasmaviruses* in *Aedes* mosquitoes from China (highlighted with green square) against other representative members in family *Phasmaviriade.* Representatives in *Orthobunyavirus* belonged to the *Peribunyaviridae* were used as the outgroup.

#### 3.3.2 Phylogeny of Phasi Charoen-like phasivirus

Since the abundance of Phasi Charoen-like phasivirus (PCLPV) reads in the *Ae. albopictus* from Guangzhou and lab-derived are relatively low, no complete genome was recovered from these pools. All three segments of PCLPV with a complete coding region were successfully recovered in one *Ae. aegypti* sample from Puerto Rico collected in 2014 and two *Ae. aegypti* samples from Kenya collected in 2018. The phylogenetic analysis was performed on these newly identified PCLPVs, all other PCLPV genomes in GenBank (including the ones identified in *Ae. aegypti* mosquitoes from Thailand in 2008, from US in 2015, from Australia in 2016, from Zhejiang Province of China in 2016, from Guadeloupe in 2017 and in Aag2 cells from UK) and two more genomes obtained from lab Aag2 cell lines in Belgium (Fig. 5). It is interesting to note that all these PCLPVs originating from distant geographic locations, different years and field mosquitoes vs. lab cell lines, all cluster very closely in the three Maximum-likelihood trees of each segment. The nucleotide similarities among the genomes of PCLPVs were very high, ranging from 94-98%, 95-99% and 94-99%, for the S, M and L segments, respectively.

**Fig. 5.**
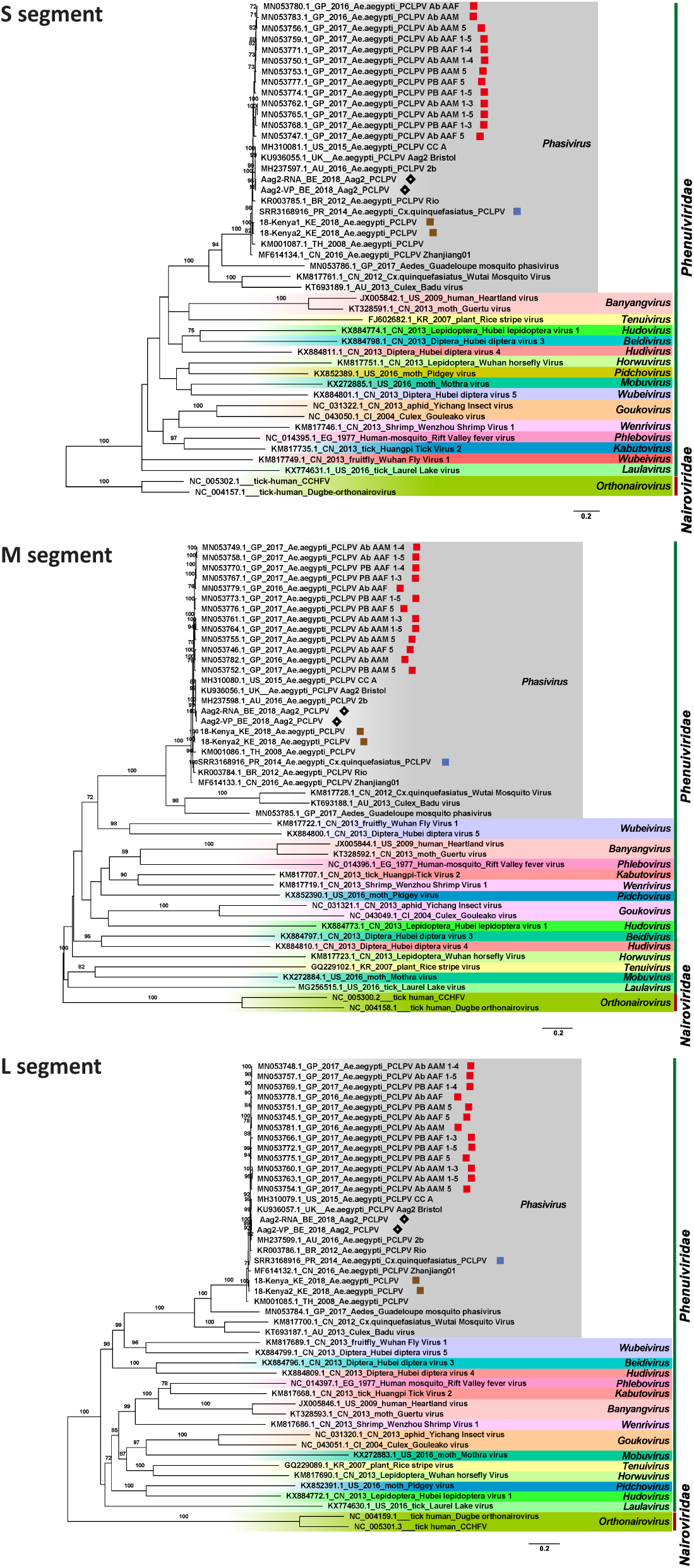
Maximum-likelihood phylogenetic tree based on nucleotide sequence of complete coding regions of the S, M, and L segment of PCLPV identified from *Aedes* mosquitoes and cells, together with other representative members of *Phenuiviriade.* The strains from Guadeloupe are highlighted with red squares, strains derived from Belgian Aag2 cells with black diamonds, strains from Kenya with brown square, and from Puerto Rico with blue square. Representative *Orthonairovirus* strains belonging to the *Nairoviridae* are used as outgroup.

#### 3.3.3 Phylogeny of Guadeloupe mosquito virus related viruses

Twelve viral genomes with complete coding regions and similar to GMV, Wenzhou sobemo-like virus 4 and Hubei mosquito virus 2 were obtained from all the analyzed virome datasets, including six pools of field *Ae. albopictus* from China (17-GZ-larvae, 17-GZ-adult, 18-GZ-larva, 18-GZ-pupa, 18-GZ-adult, SRR7204303-CN-2016), three pools of *Ae. aegypti* and *Cx. quinquefasciatus* mixture from Puerto Rico (SRR3168916-PR-2014, SRR3168921-PR-2014, SRR3168924-PR-2014), two *Ae. aegypti* pools from Kenya (18-Kenya1 and 18-Kenya2) and one from Thailand (SRR6155879-TH-2015). The viral genomes contain two segments, encoding an RdRp and one hypothetical protein on segment 1 (1500 - 1600 nt), and a capsid and one hypothetical protein encoded by segment 2 (3000 - 3200 nt). GMV, Wenzhou sobemo-like virus 4 and Hubei mosquito virus 2 are all newly described and currently unclassified viruses with distant relationship to the family *Luteoviridae* and *Sobemoviridae* [56].

The phylogenetic analysis based on segment 1 and 2 indicated that these sequences could be separated into three clades (Fig. 6). The genomes identified from Thailand in 2015, Puerto Rico in 2014 and 1 strain from Kenya clustered together with the Guadeloupe mosquito viruses found in Guadeloupe in 2016 and 2017 to form clade-1, with the nucleotide identity within this clade ranging from 90% - 99% for both segments. The genomes recovered from *Ae. albopictus* collected from Guangzhou in 2017 and 2018 cluster together with the one recovered from the same host of Yunnan in 2016 and Wenzhou sobemo-like virus 4 in clade-2, with the nucleotide identities among them ranging from 94% to 98%. The third clade consisted of the Hubei mosquito virus 2 and the sequences from 2018-Kenya2, with nucleotide identities ranging from 70% - 100%. The nucleotide similarities for both segments among these three clades ranged from 54% to 69%.

**Fig. 6.**
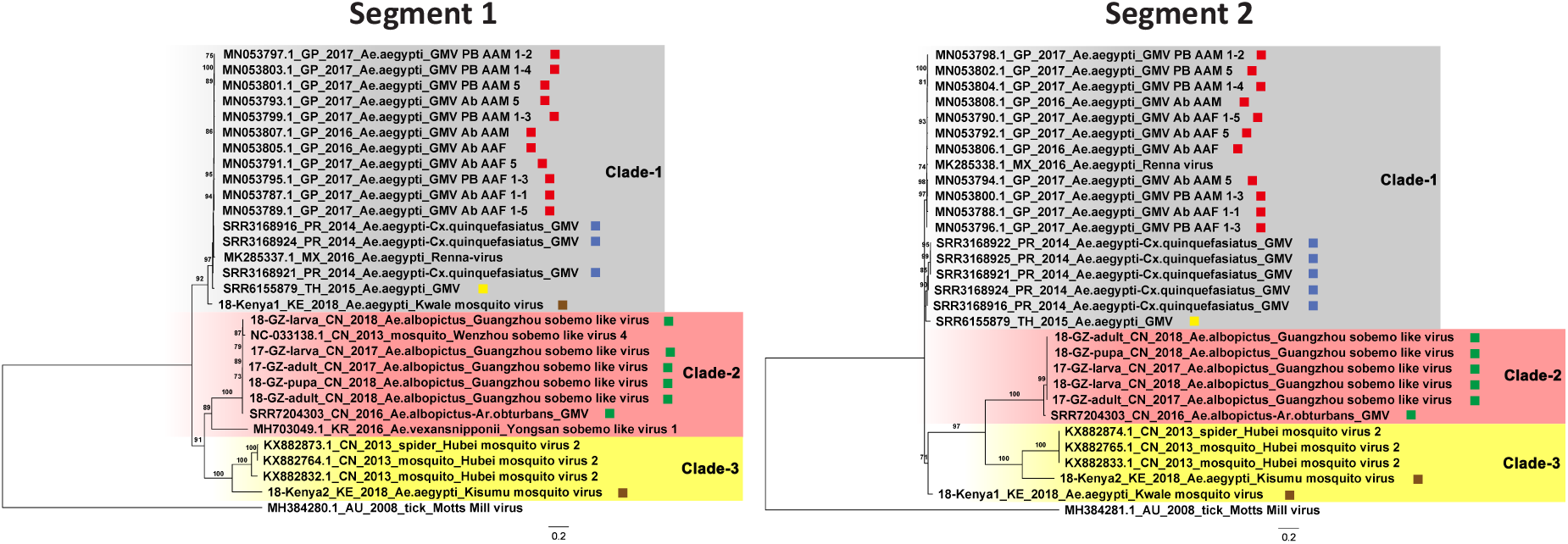
Maximum-likelihood phylogenetic tree based on nucleotide sequence of the complete coding regions in segment 1 and 2 of Guadeloupe mosquito virus related viruses. The red, blue, yellow, brown and green squares after the sequence names highlight the strains from Guadeloupe, Puerto Rico, Thailand, Kenya and China, respectively. Motts Mill virus identified from tick is used as the outgroup.

## 4. Discussion

In this study, we firstly characterized the eukaryotic virome across different developmental stages of lab-derived and field collected *Ae. albopictus* from China. As could be expected, the virome diversity in lab *Ae. albopictus* is lower than that in field mosquitoes (Fig. 2), probably due to the less complex environment and clean water and food resources. Unlike bacterial communities in mosquitoes [57–62], it is interesting to note that the virome in lab-derived *Ae. albopictus* is very stable across all stages, consistent with a vertical transmission of these viruses. Only for the larvae, relatively few viral reads were identified. This finding suggests that the virus might remain dormant at a very low concentration without or with very low rates of replication. The fact that these viruses are also present in field mosquitoes suggest that *Ae. albopictus* contains a “vertically transmitted core virome” as was described before for *Ae. aegypti* and *Culex quinquefasciatus* from Guadeloupe [56]. In addition, another set of viruses was identified to be shared by the field collected *Ae. albopictus* mosquitoes across different stages, suggesting that they have a richer “core virome”. Whether or not these additional viruses were acquired from the environment, or that the lab derived mosquitoes lost these viruses in captivity, remains to be determined. All these vertically transmitted core viromes in *Ae. albopictus* deserve more attention with respect to their effects on vector competence for important medically relevant arboviruses. The larva pool collected in the wild in 2017 is an outlier, as it contained a lot of unique viruses, which probably originated from the particular water environment they lived in, which could be contaminated by other co-inhabiting insects or larvae. Alternatively, it could be that a number of mosquitoes, other than *Ae. albopictus* were included in the pool by accident. These results further suggest a significant influence of the breeding site ecology on the mosquito viral community.

Since some viruses (e.g. PCLPV, GMV and Aedes aegypti totivirus) identified in *Ae. albopictus* from China were also identified as part of the core virome in *Ae. aegypti* from Guadeloupe [33], we were interested to find out how these core viruses in *Aedes* mosquitoes were further distributed around the world. Therefore, we explored 46 viral metagenomic datasets of *Aedes* sp. from public database (SRA) and two of *Ae. aegypti* from Kenya. Although the samples are from different countries in different continents (China, US, Australia, Kenya, Thailand, Puerto Rico, Guadeloupe), from different mosquito species (*Ae. aegypti*, *Ae. albopictus*, *Ae. alboannulatus*, *Ae. camptorhynchus*), were treated using different wet-lab and sequencing procedures, used different amounts of pooled mosquitoes and sequencing depths, a highly prevalent set of widely distributed viruses could be identified on species or family level, such as PCLPV, GMV and *Totiviridae* (Fig. 3). How these conserved viruses in *Aedes* mosquitoes interact with an arbovirus infection are interesting points for further studies. A previous study explored the effect of PCLPV together with Cell-fusing agent virus (CFAV) on the replication of arboviruses in cell line Aa23 derived from *Ae. albopictus* [63]. The Aa23 cell lines persistently infected with CFAV was further inoculated with PCLPV. CFAV-PCLPV positive Aa23 cells strongly inhibit the growth of two flaviviruses (ZIKV and DENV) and completely blocked the infection of bunyavirus La Crosse virus. Although the exact blocking mechanism is not known, this results on one hand suggest that the data generated from laboratory cell lines persistently infected by mosquito-specific viruses (MSVs) should be interpreted with caution. On the other hand, the intra-MSVs interaction need to be considered when the influence of MSVs on vector competence is explored.

Although many viruses are very prevalent in *Aedes* mosquitoes, such as Chuvirus Mos8Chu0, Kaiowa virus, Wuhan mosquito virus 6, Whidbey virus, Aedes aegypti totivirus and Australian Anopheles totivirus, phylogenetic analysis was further performed on GMV and PCLPV which had the greatest number of coding complete genomes, and APV that is highly abundant in both field and lab *Ae. albopictus* from China. APV was distantly relates to the YBV1 identified from *Ae. vexans* from South Korea forming a separate clade. All the Chinese strains from both lab and field collected mosquitoes were very similar (Fig. 4). For PCLPV, the genomes found in samples from different years, locations and habitats are very similar for all three segments (Fig. 5). Interactions between bunyaviruses and arthropods has been demonstrated over 20 million years [64–68]. The findings that both APV and PCLPV seems to be very closely related is puzzling and might suggest a rather relatively recent spread of this virus, or a very slow evolutionary rate, which might be unexpected for RNA viruses. However, the effects of these viruses on their host is very poorly understood. A recent study revealed that Aag2 cells with pre-existing infection with PCLPV do not affect the infection and growth of the mosquito-borne viruses in genus *Flavivirus*, *Alphavirus* and *Rhabdovirus* in cell culture [69]. GMV is a recently described two-segmented, currently unclassified virus. Three separate clades are observed for GMVs and GMV-related viruses, largely clustering according to locations. This group of viruses should be proposed as a new family and their role in arboviruses infection need to be further studied.

In summary, our results reveal that the virome is stable across different stages in both lab and field *Aedes albopictus*, and that a number of core viruses exists in *Aedes* mosquitoes of at least four species from seven countries. Since the number of the available *Aedes* virome data sets is still rather limited, and wet-lab procedures, sequencing depths and pool size differ strongly between the analyzed data sets, the core viruses need to be further confirmed by NGS or PCR. To fully characterize and understand the genetic and phenotypic diversity of mosquito-specific viruses, samples from individual *Aedes* mosquitoes from more locations, species and time points, processed and sequenced with the same method should be analyzed.

## Funding

This work was supported by the Ministry of Science and Technology of the People’s Republic of China [2018ZX10101004]; the National Health Commission of the People’s Republic of China [2018ZX10711001-006]; the Chinese Academy of Sciences [153211KYSB20160001]; the Wuhan Institute of Virology, China [WIV-135-PY2]; the Health Commission of Hubei Province [WJ2019Q060].

## Competing interests

The authors declare that they have no competing interests.

## Additional files

**Additional file 1:** SRA datasets used in this study

**Additional file 2:** Accession number of YBV1, PCLPV and GMV related viruses identified in this study.

**Additional file 3:** Reads number of each eukaryotic viral species in each sequencing sample

**Additional file 4:** Coverage of each eukaryotic viral species in each sequencing sample

**Additional file 5:** Alpha diversity of virome in field and lab *Aedes albopictus*

## Notes

### Competing Interest Statement

The authors have declared no competing interest.

