## Supplementary figures and images for "The conservation of a core virome in *Aedes* mosquitoes across different developmental stages and continents"

### Additional file 5

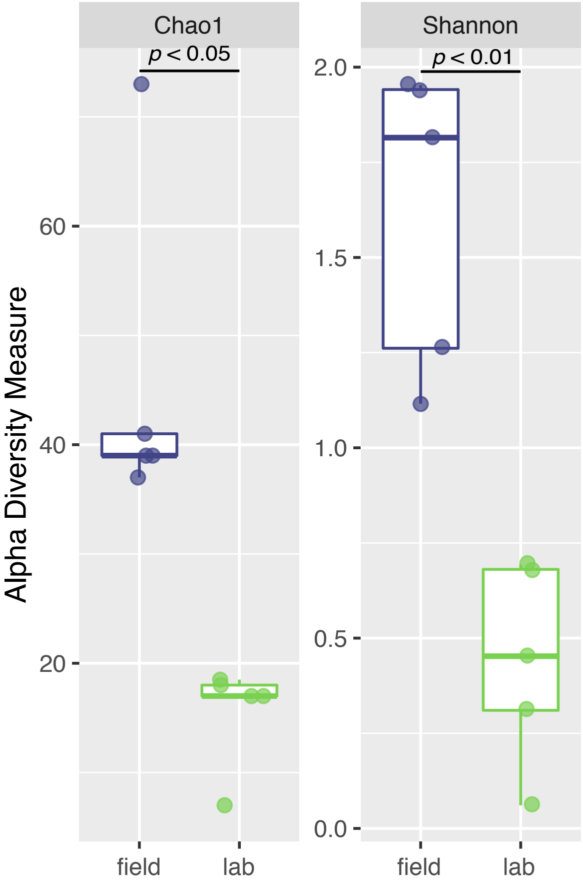
